# deMeta: Removing sub-studies from meta-analysis of genome wide association studies (GWAS)

**DOI:** 10.1101/2020.10.25.354191

**Authors:** Jiangming Sun, Yunpeng Wang

## Abstract

**Summary:** Post-GWAS studies using the results from large consortium meta-analysis often need to correctly take care of the overlapping sample issue. The gold standard approach for resolving this issue is to reperform the GWAS or meta-analysis excluding the overlapped participants. However, such approach is time-consuming and, sometimes, restricted by the available data. **deMeta** provides a user friendly and computationally efficient command-line implementation for removing the effect of a contributing sub-study to a consortium from the meta-analysis results. Only the summary statistics of the meta-analysis the sub-study to be removed are required. In addition, **deMeta** can generate contrasting Manhattan and quantile-quantile plots for users to visualize the impact of the sub-study on the meta-analysis results.

**Availability and Implementation:** The python source code, examples and documentations of **deMeta** are publicly available at https://github.com/Computational-NeuroGenetics/deMeta-beta.

**Contact:** (J. Sun); (Y. Wang)

**Supplementary information:** None.

## INTRODUCTION

Genome wide association studies (GWAS) has been very successful in discovering genomic variants that associated with human complex diseases or traits (Visscher, et al., 2017). Due to the regulations on sharing of sensitive genetic data, meta-analysis is the most deployed approach for large GWAS consortium (Willer, et al., 2010). Typically, tens of contributing groups share with the analysts of the consortium their GWAS summary statistics (sub-studies), and, the meta-analysis is performed at private computers of the consortium. Most GWAS consortiums make their meta-analysis results publicly accessible on through internet. These summary statistics is becoming an extremely valuable resources for follow-up studies on human genetic research, for example, genetic pleiotropy between diseases or traits (Bulik-Sullivan, 2015) and polygenic risk predictions (Lewis, 2020) for human complex diseases. However, these public GWAS results often include overlapping participants between one another. Not correctly accounting for the overlapping samples can generate spurious results.

Few methods have been proposed to resolve the overlapping sample issue in GWAS or post-GWAS analysis. Although the gold-standard solutions to this problem is to reperform the GWAS or meta-analysis excluding the overlapped samples, these approaches are time-consuming and sometimes not practicable due to restrictions on the data access. Lin and Sullivan proposed an approximate method (Lin and Sullivan, 2009) to correct for the overlapping samples effect in meta-analysis. And, LeBalanc et al developed a method to tackle the overlapping sample issue in genetic pleiotropic studies (LeBlanc, et al., 2018). However, no efficient program exists for removing the effect of contributing sub-studies to a consortium meta-analysis result. Under **some conditions**, exact solutions exist for resolving the overlapping issue. **deMeta** implemented the inverse of the two most popular meta-analysis frameworks (Willer, et al., 2010), *i.e.,* sample size weighted and inverse-variance weighted, in a flexible and efficient manner to fit in such situations.

## IMPLEMENTATION

We derived the inverse of sample-size weighted and inverse-variance weighted meta-analysis models in **Table 1**. The mathematical derivation is exact. We implemented the derived models into a command-line tool, **deMeta**, in Python. Our implementation has been tested successfully on three most popular operating systems, Windows 10, Linux (Ubuntu 16.04 and Red Hat Enterprise Linux 7.6), and MacOS 10.15. Other features of **deMeta** include checking and flipping DNA strands in order to match the effective SNP alleles and generating contrasting Manhattan and quantile-quantile plots (QQ -plot). The later will help the users to visualized changes in associations before and after removing overlapping samples (**Fig 1**).

**Table 1.**
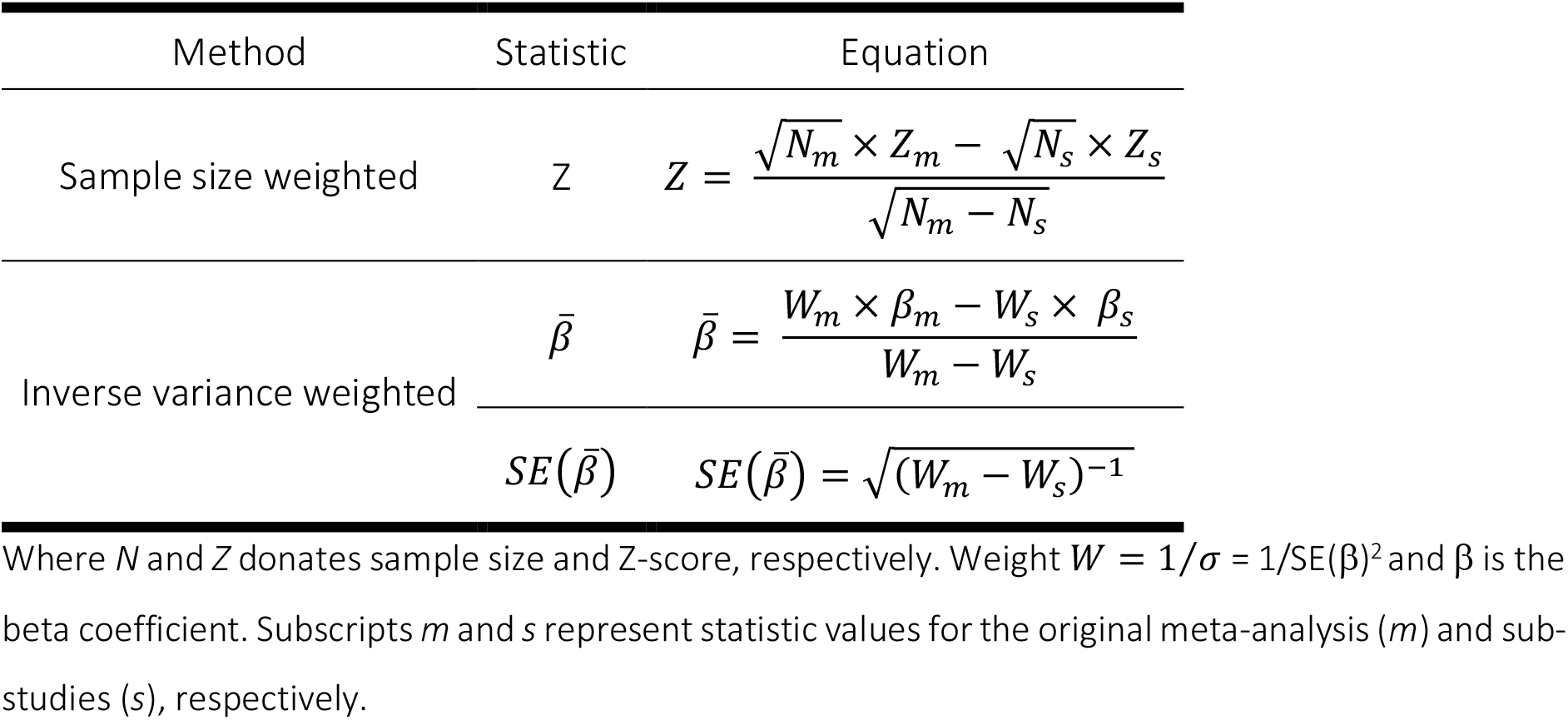
Estimator for subtracting sub-studies from a meta-analysis

| Method | Statistic | Equation |
| --- | --- | --- |
| Sample size weighted | $Z$ | $Z = \frac{\sqrt{N_m} \times Z_m - \sqrt{N_s} \times Z_s}{\sqrt{N_m - N_s}}$ |
| Inverse variance weighted | $\bar{\beta}$ | $\bar{\beta} = \frac{W_m \times \beta_m - W_s \times \beta_s}{W_m - W_s}$ |
| | $SE(\bar{\beta})$ | $SE(\bar{\beta}) = \sqrt{(W_m - W_s)^{-1}}$ |

Where $N$ and $Z$ donates sample size and Z-score, respectively. Weight $W = 1/\sigma = 1/SE(\beta)^2$ and $\beta$ is the beta coefficient. Subscripts $m$ and $s$ represent statistic values for the original meta-analysis ( $m$ ) and sub-studies ( $s$ ), respectively.

**Fig. 1.**
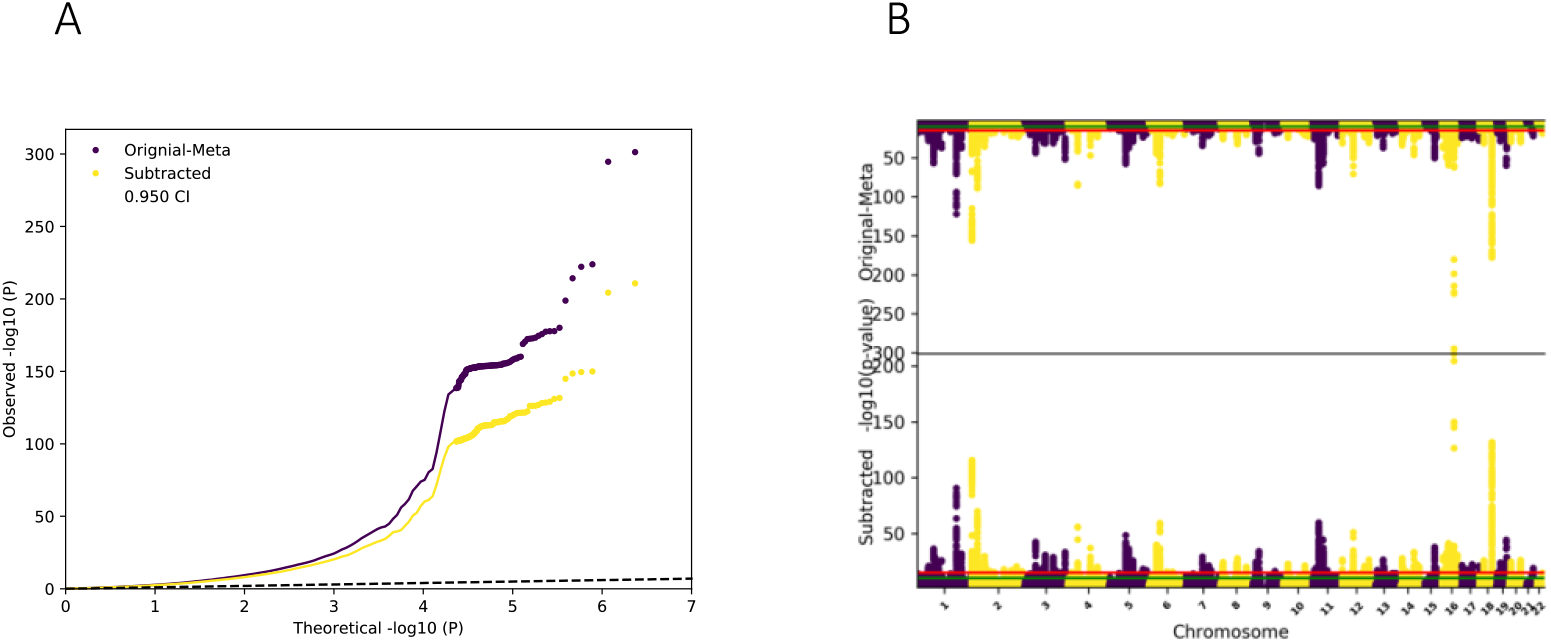
Demonstration of using **deMeta** using the GIANT BMI GWAS studies (Locke, et al., 2015; Yengo, et al., 2018). **A.** QQ plot of the GWAS summary statistics from Yengo L, *et al*. (Yengo, et al., 2018) (Original-Meta) vs after removing that from Locke AE., *et al.* (Locke, et al., 2015). **B.** Contrasting Manhattan plot for the same data sets.

## EXAMPLE

As a demonstration we applied **deMeta** to the summary statistics of the GWAS for body mass index (BMI) (Locke, et, 2015; Yengo, et al., 2018). The Yengo *et al* data is a meta-analysis results of the *Locke et al* study and the UK biobank data. Both the *Yengo et al* and the *Locke et al* data were ware downloaded from https://portals.broadinstitute.org/collaboration/giant/index.php/GIANT_consortium_data_files. The inverse of the inverse variance weighted function of **deMeta** was applied to obtain summary statistics for UK biobank data which is not available (**Fig. 1**).

## CONLCUSIONS

We have provided an efficient tool **deMeta** and demonstrated it in removing the effect of sub-studies from a meta-analysis in genetics studies. Theoretically, **deMeta** can also be used to remove effects of any subset from a large-scale consortium. Moreover, **deMeta** can eliminate/minimize effects of overlapped samples between reference and target samples when constructing polygenic risk scores for disease prediction in clinical use. Last, **deMeta** can be applied in meta-analysis beyond GWAS.

## Acknowledgements

The Computations were performed on resources provided by UNINETT Sigma2 – the National Infrastructure for High Performance Computing and Data Storage in Norway through the project NN9769K.

